# HISTONE DEACETYLASE 19 REGULATES *SHOOT MERISTEMLESS* EXPRESSION IN THE CARPEL MARGIN MERISTEM CONTRIBUTING TO OVULE NUMBER DETERMINATION AND TRANSMITTING TRACT DIFFERENTIATION

**DOI:** 10.1101/2023.05.08.537343

**Authors:** S Manrique, A Cavalleri, A Guazzotti, GH Villarino, S Simonini, A Bombarely, T Higashiyama, U Grossniklaus, C Mizzotti, AM Pereira, S Coimbra, S Sankaranarayanan, E Onelli, S Masiero, RG Franks, L Colombo

**Affiliations:** Dipartimento di Bioscienze, Università degli Studi di Milano, Via Giovanni Celoria 26, 20133, Milan, Italy; North Carolina State University, Department of Plant and Microbial Biology, Raleigh, NC, 27606, USA; Department of Plant and Microbial Biology, University of Zurich, Zollikerstrasse 107, CH-8008, Zurich, Switzerland; Institute of Transformative Bio-Molecules (ITbM), Nagoya University, Furo-cho, Chikusa-ku, Nagoya, Aichi 464-8601, Japan; Faculdade de Ciências da Universidade do Porto, Departamento de Biologia, Universidade do Porto, rua do Campo Alegre, 4169-007 Porto, Portugal; LAQV Requimte, Sustainable Chemistry, Universidade do Porto, 4169-007 Porto, Portugal; Department of Biological Sciences and Engineering, Indian Institute of Technology Gandhinagar, Palaj, Gujarat 382355, India

## Abstract

The gynoecium is critical for the reproduction of flowering species as it contains the ovules and the tissues required for pollen germination and guidance. These tissues are collectively known as the reproductive tract (ReT) and comprise stigma, style and transmitting tract (TT). The ovules and the ReT originate from a meristem within the pistil named carpel margin meristem (CMM).

SHOOT MERISTEMLESS (STM) is a key transcription factor required for meristem formation and maintenance. In all above-ground meristems, including the CMM, *STM* has to be locally downregulated to allow proper organ differentiation. However, how this downregulation is achieved in the CMM is unknown.

In this work, we have studied HISTONE DEACETYLASE 19 (HDA19) role in ovule and ReT differentiation, based on the observation that *hda19-3* mutant displays reduced ovule number and fails to properly differentiate the TT. Fluorescence activated cell sorting (FACS) coupled with RNA-seq revealed that in the CMM of *hda19-3* mutant, genes promoting organ development are downregulated while meristematic markers, including *STM*, are upregulated. We found that HDA19 is fundamental to downregulate *STM* in the CMM, allowing ovule formation and TT differentiation. *STM* is ectopically expressed in *hda19-3* at intermediate stages of pistil development, and its downregulation by RNA interference alleviated *hda19*-3 phenotypic defects. Furthermore, chromatin immunoprecipitation assays indicated that *STM* is a direct target of HDA19 during pistil development and that *SEEDSTICK* (*STK)* is required for the histone acetylation-mediated regulation of *STM*.

Our results have led to the identification of the factors required for *STM* silencing in the gynoecium allowing organogenesis and tissue differentiation from the CMM.

## Introduction

The number of seeds produced by a plant is important for the reproductive success of species and for food security. To increase seed number, plants can produce more flowers and/or more seeds per flower. As seeds are formed upon ovule fertilization, final seed number depends partially on ovule number.

The ovules and the reproductive tract (ReT) emerge from the carpel margin meristem (CMM) (Reyes-Olalde et al., 2013a; Chávez Montes et al., 2015). The CMM is a determinate meristem that appears as a ridge in the medial part of the gynoecium at stage 7 of floral development (Figure 1A and Smyth et al., 1990). At stage 8, it produces the placenta, from which ovule primordia (OP) arise at stage 9. Contemporarily to OP formation, the medial ridges fuse forming the septum (Figure 1A). From stages 10 to 12, ovules, stigma, style and transmitting tract (TT) complete their developmental program, and the gynoecium becomes competent for fertilization (Bowman et al., 1999; Roeder and Yanofsky, 2006; Reyes-Olalde et al., 2013b; Chávez Montes et al., 2015; Herrera-Ubaldo et al., 2019).

**Figure 1.**
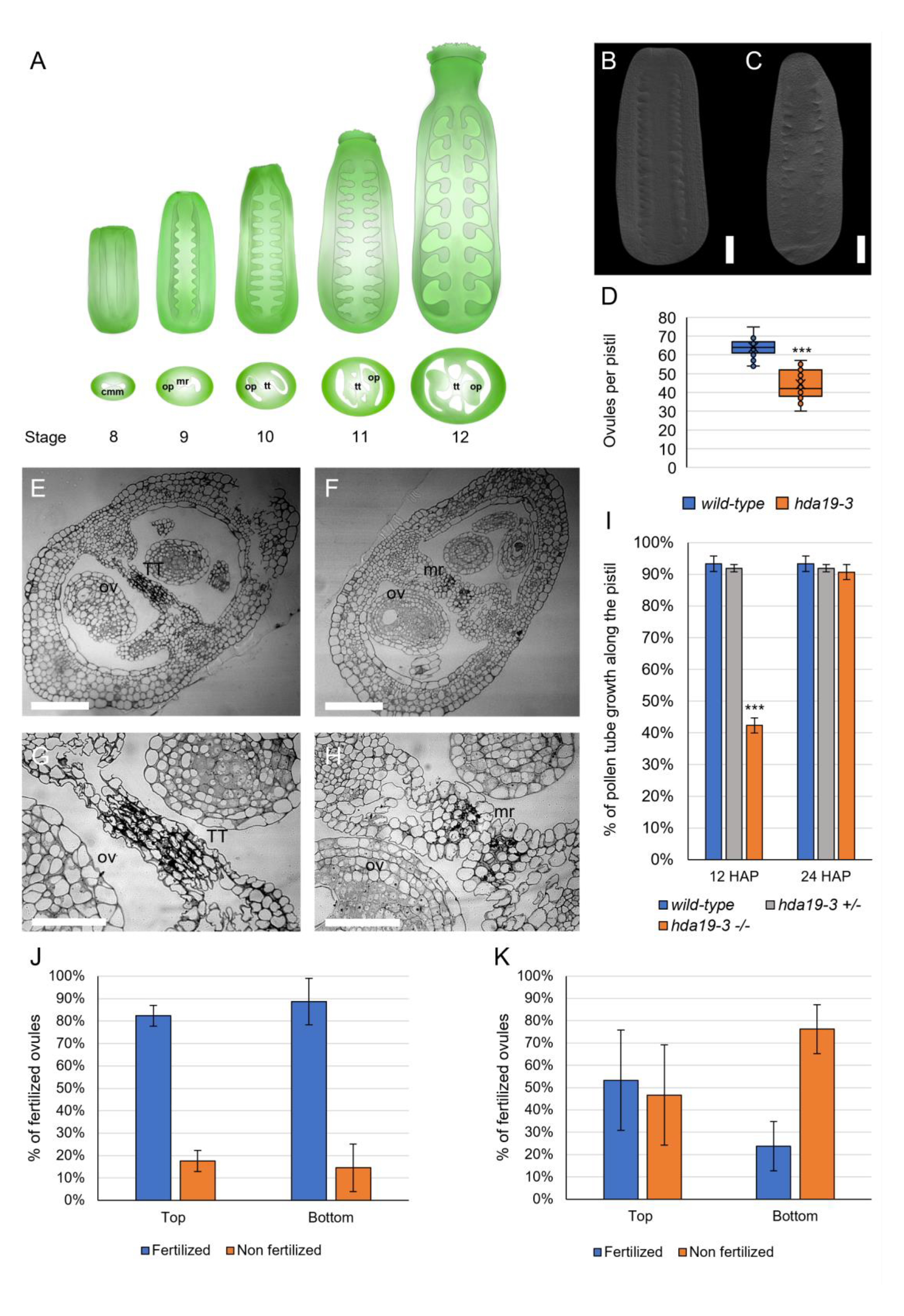
Ovule density and transmitting tract are abnormal in *hda19-3*. **A)** Scheme of the development of the reproductive tract from the CMM from stage 8 to 12 of flower development according to Smyth et al., 1990. Abbreviations: CMM=carpel margin meristem; OP=ovule primordia; TT= transmitting tract. **B-C)** Cleared pistils of *wild-type* (**B**) and *hda19-3* (**C**) at stage 9, showing differences in ovule density. Scale bar = 50 µm. **D)** Ovule number of *wild-type* (n = 70 pistils) and *hda19-3* (n= 85 pistils) plants. Bars represent the mean ± s.e. value. *** = Student’s T-test with p-value < 0.001. **E-H)** Transversal section of *wild-type* (**E**, **G**) and *hda19-3* (**F**, **H**) pistils at stage 12. *hda19-3* shows unfused septum and defective transmitting tract due to the lack of degeneration of septum cells. Images **G** and **H** are close up images of a portion of images **E** and **F** respectively. Abbreviations: TT = transmitting tract; OV = ovule; mr = medial ridge. Scale bar = 50 µm. **I)** Ratio of pollen tube length/pistil length at 12 and 24 HAP in *wild-type* (n= 8), heterozygous (n= 8) and homozygous (n= 12) *hda19-3* pistils, hand-pollinated with wild-type pollen. Bars represent the mean ± s.e. value. *** = Student’s T-test with p-value < 0.001. **J-K)** Seed set at the top-half and bottom-half of *wild-type* (n = 10) (**J**) and *hda19-3* (n = 10) (**K**) siliques, hand-pollinated with *wild-type* pollen, showing that the bottom part of *hda19-3* siliques shows higher levels of unfertilized ovules than the top. Bars represent the mean ± s.e. value.

Ovule formation encompasses two steps: (1) boundary formation and primordium outgrowth and (2) identity determination. The first step is controlled by a network of genes that modulate lateral organ formation such as CUP-SHAPED COTYLEDON 1/2/3 (CUC1/2/3), BLADE-ON-PETIOLE (BOP), PIN-FORMED 1 (PIN1), and AINTEGUMENTA (ANT) (Ishida et al., 2000; Mizukami and Fischer, 2000; Vernoux et al., 2000; Norberg et al., 2005). By contrast, the second step is controlled by organ-specific gene networks. Ovule identity is redundantly determined by three MADS-box transcription factors: SEEDSTICK (STK) and SHATTERPROOF1/2 (SHP1/2). In the triple mutant *stk shp1 shp2*, ovules are replaced by carpelloid structures (Favaro et al., 2003; Pinyopich et al., 2003).

As for ReT tissues, the TT supports the growth of pollen tubes to reach the ovules. It forms in the center of the septum and runs from the top to the base of the pistil (Figure 1A) (Crawford and Yanofsky, 2008). In *Arabidopsis*, pollen tubes grow through the intercellular spaces between TT cells, and their growth is aided by the production of a facilitating extra-cellular matrix, and by the death of the TT cells. This process is developmentally regulated and starts before anthesis (Crawford and Yanofsky, 2008; Pereira et al., 2021). The formation of the TT initially depends on transcription factors controlling the formation of the septum like ETTIN (ETT), SPATULA (SPT), SEUSS (SEU), ANT or INDEHISCENT (IND) (Alvarez & Smyth, 2002; Azhakanandam et al., 2008; Kay et al., 2013; Pereira et al., 2021; Sessions & Zambryski, 1995). Later, genes like *NO TRANSMITTING TRACT* (*NTT*) and *HECATE1/2/3* (*HEC1/2/3*) control the maturation of TT cells (Gremski et al., 2007, Herrera-Ubaldo et al., 2019; Marsch-Martínez et al., 2014). Interestingly, STK and SHP1/2 are also required for TT formation in cooperation with NTT (Marsch-Martínez et al., 2014; Herrera-Ubaldo et al., 2019) and CESTA/HALF-FILLED (CES) (Di Marzo et al., 2020).

The CMM is a determinate meristem, so that stem cells are maintained transiently, and consumed once ovule and tissue differentiation are completed (Alvarez & Smyth, 2002; Reyes-Olalde & de Folter, 2019). Nevertheless, genetic and transcriptomic analyses have shown its similarity with undetermined meristems like the shoot apical meristem (SAM) (Wynn et al., 2011; Heidstra and Sabatini, 2014; Gaillochet and Lohmann, 2015; Villarino et al., 2016; Reyes-Olalde and de Folter, 2019; Zúñiga-Mayo et al., 2019).

HISTONE DEACETYLASE 19 (HDA19) is a major regulator of the maintenance of the SAM (Long et al., 2006; Gorham et al., 2018; Chung et al., 2019a), the flower meristem (FM) (Krogan et al., 2012a; Bollier et al., 2018) and the root apical meristem (RAM) (Pi et al., 2015a). HDA19 is an epigenetic modifier that removes acetyl groups from histones, repressing the transcription of the genes associated with such histones (Kumar et al., 2021). HDA19 target specificity is provided by the interaction with transcription factors and other cofactors (Kumar et al., 2021). For instance, in the inflorescence meristem (IM), a complex between HDA19 and the auxin response factors ETT, ARF4 and the YABBY FILAMENTOUS FLOWERS (FIL) downregulates the *KNOTTED-1-like* homeobox (*KNOX*) transcription factor *SHOOT MERISTEMLESS (STM*), allowing flower primordia specification (Chung et al., 2019). Interestingly, it has been previously suggested that STM role is similar in both the SAM and the CMM (Scofield et al., 2007; Durbak and Tax, 2011; Poulios and Vlachonasios, 2018) as it is required to maintain meristematic identity in both contexts. *STM* has to be locally downregulated to allow the formation of organ primordia from the SAM (Long et al., 1996; Kanrar et al., 2006; Landrein et al., 2015). Likewise, in pistils, its overexpression interferes with ovule identity determination and septum formation (Scofield et al., 2007).

Here, we show that mutation of *HDA19* leads to ovule number reduction and TT defects. We identified targets of HDA19 during TT differentiation and ovule formation, through transcriptomic analysis of cells isolated by fluorescence activated cell sorting (FACS), using a *STK* marker line active in the TT, placenta, and ovules. We observed by RNA-seq and in situ hybridization (ISH) that *STM* was over-and ectopically expressed in TT and ovules in *hda19-3* in respect to *wild-type*. Furthermore, chromatin immunoprecipitation (ChIP) experiments showed that *STM* locus was more acetylated in *hda19-3* flowers. Our work shows also that *stk* mutant has similar defects to *hda19-3* in ovule number and TT differentiation and that STK binds *STM* locus and regulates its acetylation state. Overall, our results indicate that correct development of ovules and TT requires the repression of *STM* by HDA19 and that STK is also necessary for histone acetylation-mediated downregulation of *STM* in the CMM.

## RESULTS

### Development of the reproductive tract requires HDA19

It was previously observed that *hda19* mutants have a reduced seed set (Zhou et al., 2013), a phenotype that can be caused by pre- (Kuhn et al., 2020) and/or post-fertilization defects (Zhou et al., 2013). Despite these descriptions, the molecular basis of *hda19* reduced seed production has not been investigated. Therefore, in this work we have focused on the involvement of HDA19 on pre-fertilization stages and specially in its role during ovule number determination and TT differentiation.

We performed a phenotypic characterization on *hda19-3* mutant pistils considering traits as ovule density and reproductive tract morphology. At first instance, we analysed ovule number in *wild-type* and *had19-3*. We observed that, even though *hda19-3* pistils have similar size respect to *wild-type* (Figure 1B, C), they contain significantly fewer ovules (51±3) compared to *wild-type* (67±4 ovules) (Figure 1D), suggesting that ovule density is lower. Next, we analysed pistils radial organization observing that, in *hda19-3,* the medial ridges remain unfused at stage 12 (Figure 1E-H). Consequently, the TT is not properly formed and the degeneration of the subepidermal cells required for TT maturation is also drastically reduced in *hda19-3* (Figure 1F, H) respect to the *wild-type* (Figure 1E, G). These defects impair pollen tube growth along the pistil (Figure 1I, Supplemental Figure 1), and reduce fertility predominantly at the bottom-half of the silique (Figure 1J, K). In hand-pollination experiments using *wild-type* pollen, the percentage of unfertilized ovules in *wild-type* siliques at 16 hours after pollination was low and similar between the top-and bottom -half of the siliques (Figure 1J). In *hda19-3*, fertility is overall reduced in respect to *wild-type* (Figure 1J, K). In addition, the percentage of unfertilized ovules at the bottom-half of the siliques was significatively higher respect to the top-half (Figure 1K). Altogether, we concluded that HDA19 is required for proper ovule primordia specification and TT development.

### Transcriptomic analysis of *hda19-3* sorted cells supports a role of HDA19 in reproductive tract development and ovule number determination

As *HDA19* is highly expressed in most tissues of the inflorescence (Zhou et al., 2005) and we wanted to focus on its role during ovule number determination and TT differentiation, we designed a strategy based on the isolation of these specific cell types to perform transcriptomic analysis.

We crossed *hda19-3* with a line containing the genomic region of *STK* fused to *GFP* (*pSTK::STK-GFP,* Mizzotti et al., 2014), as *STK* is expressed in the placenta, ovules and TT from stage 8 to 12 (Mizzotti et al., 2014; Herrera-Ubaldo et al., 2019). After verifying that the spatial expression pattern of *pSTK::STK-GFP* was equivalent in both genetic backgrounds (Supplemental Figure 2), fluorescent protoplasts were isolated by FACS and RNA was extracted for RNA-sequencing to identify differentially expressed genes.

Before proceeding with library preparation, enrichment of *STK-GFP* in sorted protoplasts was validated by qRT-PCR (Supplemental Figure 3). After the validation, three samples of RNA from *wild-type* GFP-positive cells, and four samples from *hda19-3* GFP-positive cells (80-100.000 cells/replicate) were used for library preparation and RNA Illumina sequencing (HiSeq2500).

Since HDA19 is an epigenetic regulator, we used a stringent approach to narrow down the number of differentially expressed genes (DEGs). We used Cufflinks (Trapnell et al., 2012), DeSeq2 (Love et al., 2014) and EdgeR (Robinson et al., 2009) to analyze the RNA-seq data (Supplemental File 1-3). Then, we compared the DEGs obtained by the three software and selected only those in common among the three programs (Supplemental File 4). This approach retrieved 554 upregulated genes and 408 downregulated genes (Figure 2A).

**Figure 2.**
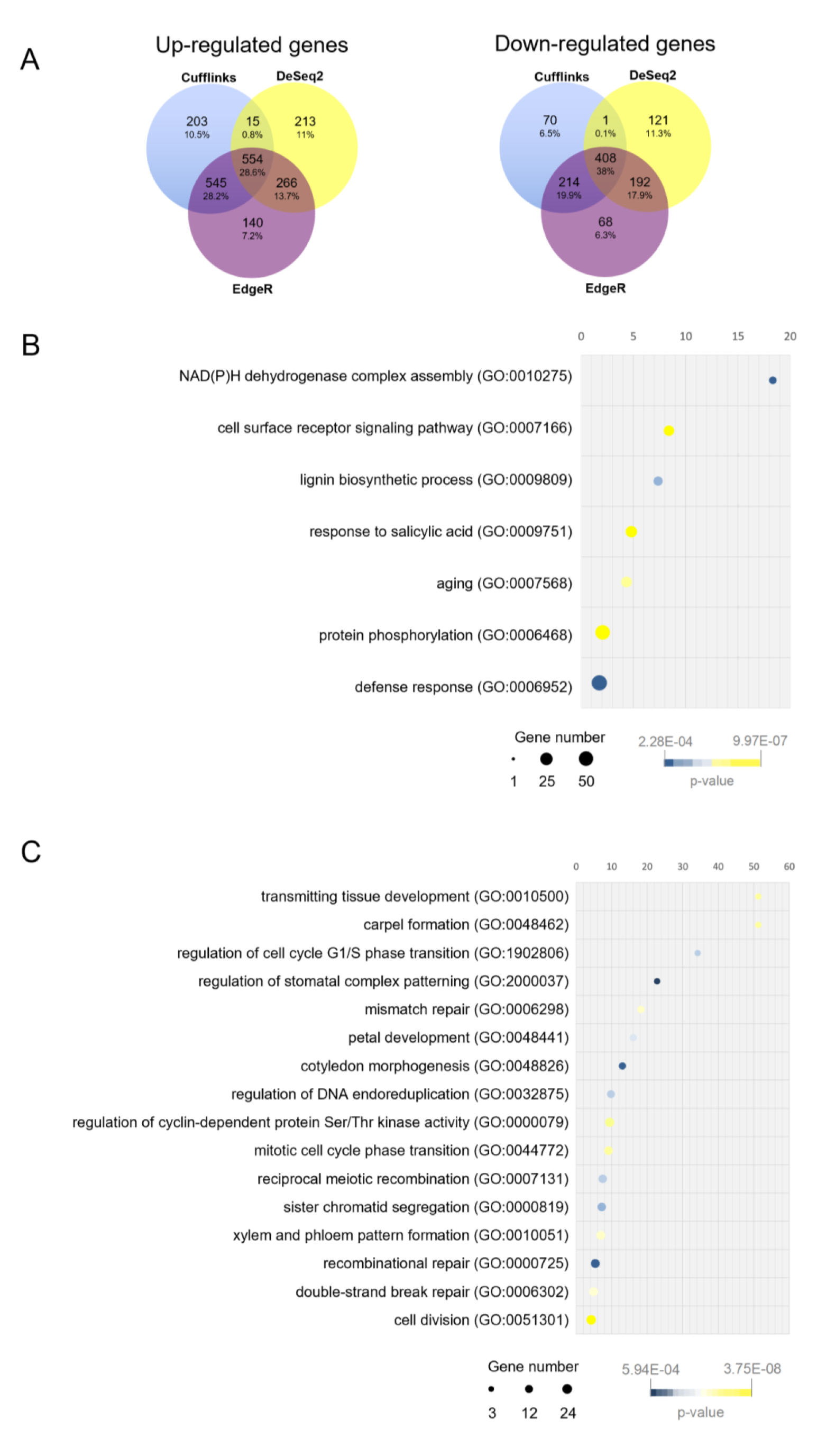
Reproductive tract transcriptomic comparison of *wild-type* and *hda19-3.* **A)** Venn diagrams showing the overlap of up-and down-regulated genes (FDR<0.001 and logFC>|1.5|) detected by Cufflinks, EdgeR and DeSeq2. **B)** Enrichment analysis of ‘Biological Process’ GO terms associated with up-regulated genes (logFC>|1.5|) detected by all three programs. **C)** Enrichment analysis of ‘Biological Process’ GO terms associated with down-regulated genes (logFC>|1.5|) detected by all three programs (fold enrichment > 4). Graphs show overrepresented child-most (tip) GO terms. Parent GO terms (root terms) have not been plotted for simplicity. Enrichment analysis was performed using Panther (Mi et al., 2018, 2019) with the default parameters set by the program.

To survey the roles of the DEGs, we performed an enrichment analysis of ‘Biological Process’ GO terms associated with the upregulated and downregulated genes (Figure 2 B, C and Supplemental File 5, 6) using Panther (Mi et al., 2018, 2019). Among the upregulated genes we found GO terms as ‘response to salicylic acid’, ‘defense response’ or ‘aging’ that directly reflect previously described roles of HDA19 (Zhou et al., 2005; Kim et al., 2008; Tanaka et al., 2008; Zhu et al., 2010; Choi et al., 2012) (Figure 2B, Supplemental File 5). Likewise, ‘NAD(P)H dehydrogenase complex assembly’ relates to the role of HDA19 in light response regulation (Benhamed et al., 2006; Guo et al., 2008; Jan et al., 2011).

Among the downregulated genes, the most enriched GO terms (Figure 2C) were ‘transmitting tissue development’ and ‘carpel formation’, both with a fold enrichment of 51.28 (Figure 2C, Supplemental File 6).

Overall, GO terms enrichment analysis showed that genes associated with known roles of HDA19 were present among the DEGs, supporting the validity of our dataset and of the sampling strategy, and revealed that genes necessary for the normal progression of carpel and TT development were downregulated in *hda19-3* sorted cells.

### Meristematic markers and genes involved in ovule and transmitting tract development are deregulated in *hda19-3* mutant

HDA19 regulates the maintenance of the stem cell population in several meristems (Long et al., 2006; Krogan et al., 2012b; Pi et al., 2015b; Bollier et al., 2018). Therefore, we checked for deregulated meristematic markers in our dataset, and we found that *CLAVATA1* (*CLV1*) as well as the KNOX genes *STM*, *KNAT1/BREVIPEDICELLUS* (*BP*) and *KNAT 6* showed higher expression in *hda19-3* in respect to *wild-type* (Figure 3A).

**Figure 3.**
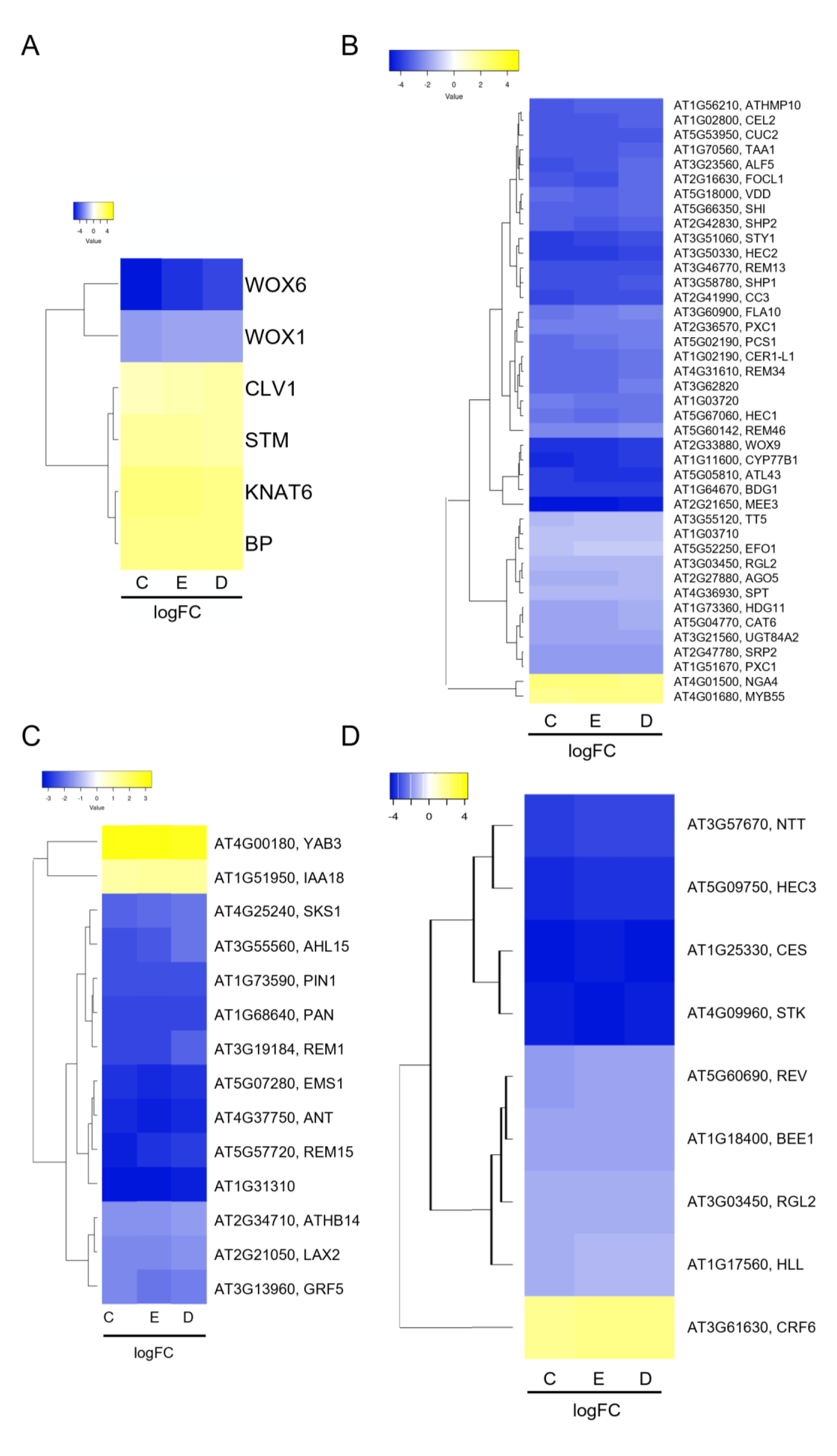
CMM-specific genes are deregulated in *hda19-3* mutant. **A)** Meristem marker genes deregulated in *hda19-3* mutant. **B)** CMM-specific genes from stages 6-8 of pistil development according to Villarino *et al*. (2016) that are deregulated in *hda19-3*. **C)** CMM-specific genes from stages 8-10 of pistil development according to Wynn *et al*. (2011) that are deregulated in *hda19-3.* **D)** Notable ovule number and TT regulators are deregulated in *hda19-3*. Genes plotted correspond to genes with an FDR < 0.001 and logFC > |1.2| in the RNA-seq data according to Cufflinks (C), EdgeR (E) and DeSeq2 (D). Genes were hierarchically clustered by average clustering using Euclidean distances. Heatmaps were obtained using Heatmapper (http://heatmapper.ca/). The color scale represents the logFC (*wild-type* vs *hda19-3*) as detected by each program and the color is relative to the maximum and minimum values of each table.

*STM*, *BP* and *KNAT6* are partially redundant for SAM maintenance (Byrne et al., 2002), and their local downregulation at the boundaries of the meristem is necessary for organ initiation (Belles-Boix et al., 2006; Ragni et al., 2008; Zhao et al., 2015). Therefore, their upregulation in *hda19-3* could be hindering the formation of CMM-derived structures.

Furthermore, to identify CMM-specific genes associated to ReT differentiation and ovule initiation, we cross-referenced our datasets with data from two articles that described sets of CMM-specific genes (Wynn et al. 2011; Villarino et al., 2016). Regarding Villarinós set (stages 5-8), we found that several genes involved in TT and style development like *HEC1*, *HEC2*, *SPT* (Gremski et al., 2007; Nahar et al., 2012) and *STYLISH* (*STY1*) (Sohlberg et al., 2006; Kuhn et al., 2020) were downregulated in *hda19-3* (Figure 3B). This might explain the defects in the TT (Figure 1E-H) and the style (Kuhn et al., 2020) of *hda19-3.* Additionally, several well-known players of ovule initiation from Wynn’s dataset (stages 8-10) (Wynn et al., 2011a), like *PHABULOSA* (*PHB*), *ANT*, *PERIANTHIA* (*PAN*) and *PIN1* (Azhakanandam et al., 2008; Cucinotta et al., 2014; Galbiati et al., 2013; Lee & Clark, 2015; Nole-Wilson, 2006) were downregulated in *hda19-3* (Figure 3C). This observation is consistent with the reduced number of ovules observed in *hda19-3* (Figure 1D).

Only four CMM-specific genes were upregulated in *hda19-3*: *NGATHA4* (*NGA4*) and *MYB55* (Villarino et al., 2016) (Figure 3B); and *YABBY3* (*YAB3*) and *INDOLE-3-ACETIC ACID INDUCIBLE 18 (IAA18)* (Wynn et al., 2011) (Figure 3C). *NGA4* and *YAB3* are negative regulators of lateral organ formation (Lee et al., 2015, Alvarez et al., 2009, González-Reig et al., 2012), agreeing with *hda19- 3*’s ReT phenotypes. The role of *IAA18* and *MYB55* in the pistil is not known, but *iaa18* mutants have reduced fertility (Uehara et al., 2008), and *MYB55* is a direct target of BRASSINAZOLE-RESISTANT 1 (BZR1) (He, 2005), which participates in organ boundary establishment in the SAM (Bell et al., 2012; Gendron et al., 2012). Therefore, upregulation of these genes might also interfere with ReT development.

Finally, we checked the expression of some additional regulators of TT and ovule number determination that were not included in Villarino’s or Wynn’s datasets due to the experimental conditions chosen in their work. For example, *HEC3* and *STK* itself were downregulated (Figure 3D). It is worth noting that, although *STK* shows a logFC close to −4 in *hda19-3*, STK spatial pattern in *hda19-3* was unchanged in respect to *wild-type* (Supplemental Figure 2).

Overall, analysis of CMM-specific genes revealed that several *KNOX* genes involved in meristem maintenance (*STM*, *BP*, *KNAT6*) and negative regulators of lateral organ development (*NGA4* and *YAB3*) were upregulated in *hda19-3* pistils, while positive regulators of TT differentiation and ovule initiation were downregulated (e.g., *PIN 1* and *ANT*). As HDA19 is a transcriptional repressor, this suggests that HDA19 might directly regulate meristematic genes and/or genes involved in repressing the proper differentiation and formation of CMM-derived structures.

### *STM* is ectopically expressed and over-acetylated in the reproductive tract of *hda19-3* mutant

Based on its role in other meristems and on our data, we hypothesized that *STM* could be one of the main end-goal of HDA19’s repressing activity in the CMM. To gain insight on the spatial expression pattern of *STM* in *wild-type* and *hda19-3* pistils, we performed *in-situ* hybridization (ISH) with a *STM*-specific antisense probe (Figure 4A-H). At stage 7, the expression pattern of *STM* was similar in both genotypes (Figure 4A, E). However, from stages 8 to 10, the expression of *STM* in the medial ridge increased in *hda19-3* and expanded to the placenta, OP and even to the lateral domain (Figure 4B-C, F-G). After stage 10, *STM* expression became restricted to the adaxial replum zone in both genotypes (Figure 4D, H). As STM promotes meristematic fate, its ectopic expression in the placenta, OP and TT from stage 8-10 might interfere with ovule initiation and TT development, causing the phenotypes observed in *hda19-3* mutant.

**Figure 4.**
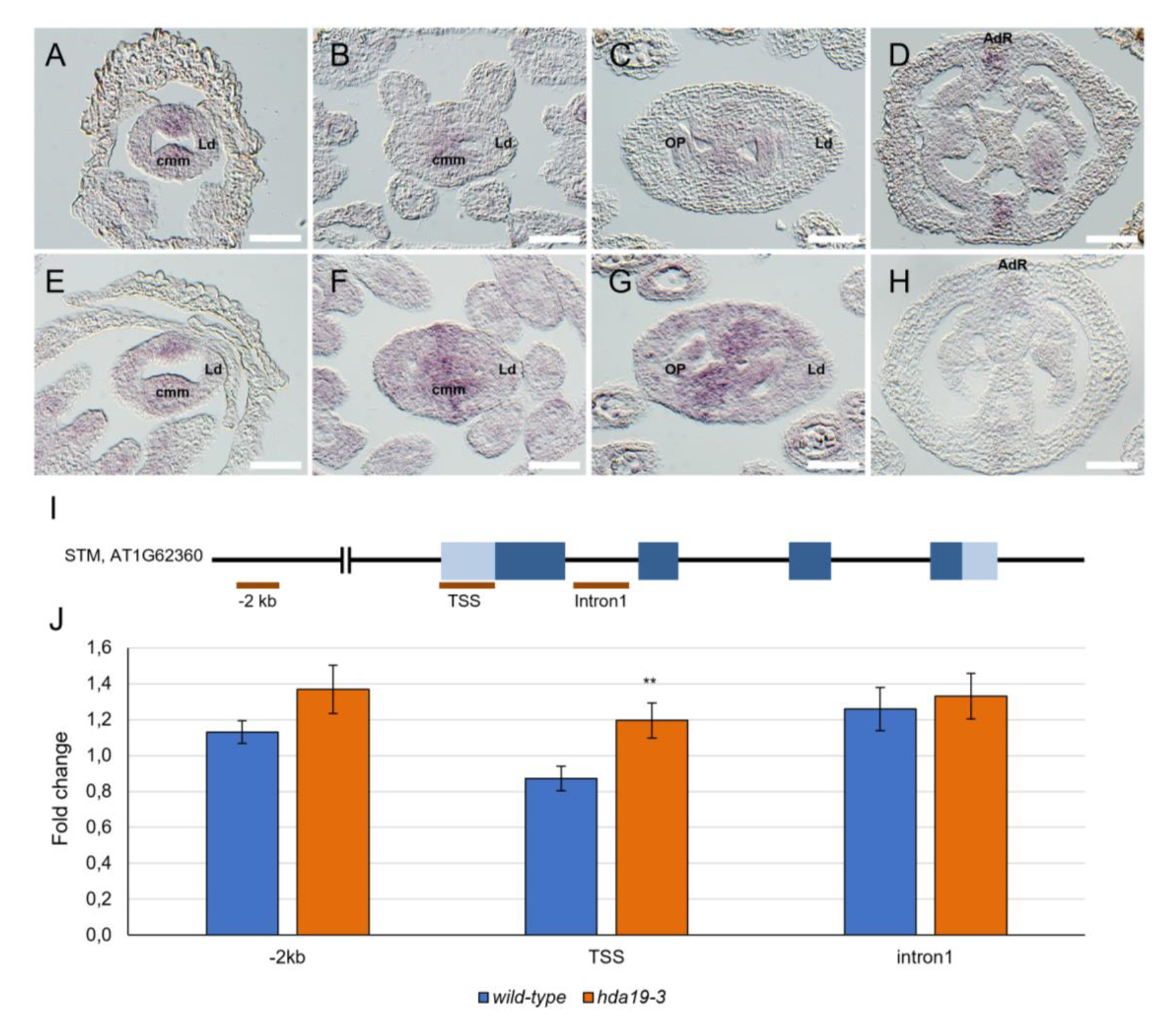
*STM* is ectopically expressed and over-acetylated in *hda19-3* pistils. **A-H)** Detection of *STM* mRNA by *in-situ* hybridization in *wild-type* **(A-D)** and *hda19-3* **(E-H)** pistils. At stage 6, *STM* mRNA is specifically expressed in the CMM of *wild-type* pistils (**A-B**). Then expression decreases along following stages until getting restricted to the adaxial replum zone (**C-D**). Instead, in *hda19-3* during stages 7-10, when ovule primordia arise, the expression of *STM* increases and expands outside of the CMM (**F-G**), to finally become also restricted to the replum later on (**H**). Abbreviations: CMM = carpel margin meristem; LD= lateral domain; OP = ovule primordia; AdR = adaxial replum. Scale bar = 50 µm. **I)** Regions of *STM* locus tested for changes in AcH3K9/14. **J)** AcH3K9 ChIP-qPCR of *STM* locus in *wild-type* and *hda19-3*. Graph shows the mean ± sd result of *STM* regions enrichment in *hda19-3* background in respect to the *wild-type*, among three independent biological replicates of the AcH3K9 ChIP-qPCR experiment performed on *wild-type*, *hda19-3, stk* and *stkshp1shp2* flowers. ** = Student’s T-test with p-value < 0. 01.

To analyze whether the changes in *STM* expression pattern could be caused by variations in its histone acetylation levels, resulting from the absence of HDA19, we collected flowers at stages 1-12 from *wild-type* and *hda19-3* plants and performed ChIP followed by quantitative Real-Time PCR (ChIP-qPCR). We used an antibody against HISTONE 3 (H3) acetylated in Lysine 9 and 14 (K9/14) (AcH3K9/14) (Figure 4I-J). Since no other histone de/acetylases are significantly deregulated in *hda19-3* sorted cells (Supplemental Figure 4), the changes in *STM* histones acetylation observed with this antibody should be caused mostly by the direct action of HDA19.

We tested the acetylation levels in several regions of *STM* locus: (1) the distal promoter (−2 kb) (Figure 4I), because it was reported that HDA19 can deacetylate *STM* in these regions (Chung et al., 2019a); (2) the transcription start-site (TSS), as it generally correlates with active gene expression (Wang et al., 2017; Kim et al., 2021; Kumar et al., 2021); and (3) the first intron. We observed a significant increase in acetylation at the *STM* TSS in *hda19-3*, and a moderate increase at the promoter region, while no significative increment in acetylation was found in the first intron (Figure 4J). In conclusion, our results suggest that at the stages tested, HDA19 is required for the deacetylation of *STM*.

### Downregulation of *STM* alleviates reproductive tract defects of *hda19-3* mutant in an expression-level dependent manner

To verify if the ectopic expression of *STM* was involved in the reduction of ovule number and in the TT defects of *hda19-3*, we used an RNA interference (*RNAi*) construct to reduce *STM* levels in *hda19-3* mutant background. (Fornara et al., 2004).

We analyzed eight *hda19-3* independent lines transformed with *35S::STM-RNAi::pFRH* (Figure 5, Supplemental Figure 5). We measured *STM* mRNA levels in the inflorescences of *wild-type*, *hda19-3* and *35S::STM-RNAi::pFRH* (*hda19-3 STMRNAi* hereafter) transformed plants by qRT-PCR. The levels of *STM* transcript in *hda19-3* mutant were 2-fold those of *wild-type* (Figure 5A). All the analyzed *hda19-3 STMRNAi* transformants showed levels of *STM* mRNA in between those of *hda19-3* and the *wild-type* (Figure 5A).

**Figure 5.**
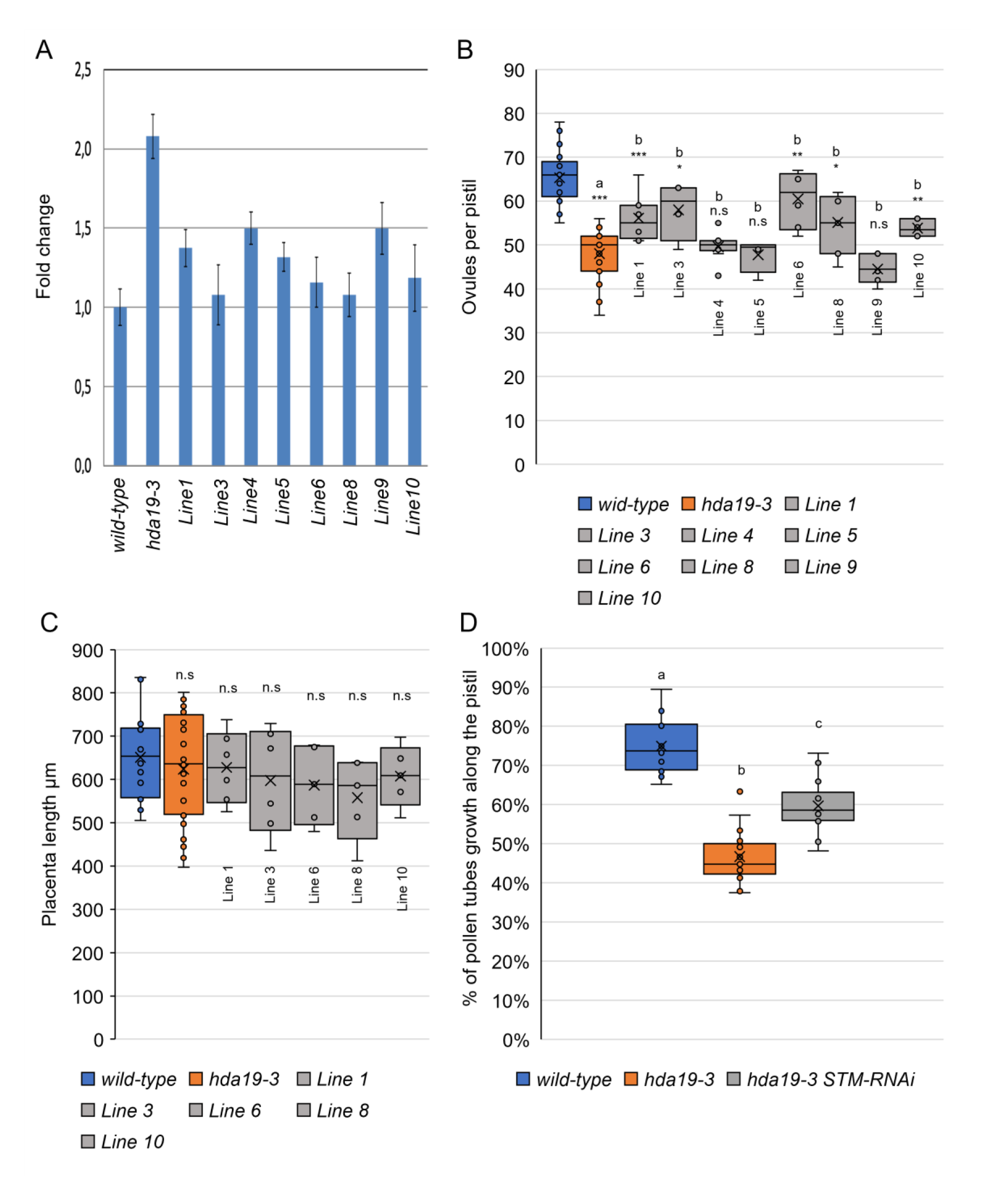
Downregulation of *STM* alleviates *hda19-3* reproductive tract phenotypes. **A)** Level of *STM* mRNA in inflorescences of *wild-type*, *hda19-3* and 8 independent *hda19-3 STMRNAi* lines. **B)** Ovule number per pistil of *wild-type* (n=25), *hda19-3* (n=27) and *hda19-3 STMRNAi* lines (in order from line 1 to line 10: n=8, 4, 10, 4, 6, 7, 6, 6). The statistical significance of ovule number was determined by ANOVA and a Bonferroni *post-hoc* test for multiple comparisons. Letters over boxes denote significantly different averages and asterisks the level of signification (* = p-value < 0.05; ** = p-value < 0.01; *** = p-value < 0.001). **C)** Placenta length of *wild-type*, *hda19-3* and the *hda19-3 STMRNAi* lines that showed significant changes in ovule number (Lines 1, 3,6, 8, 10). **D)** Pollen tube growth measured as pollen tube length /total length of the pistil. Pollen tubes were allowed to grow for 14h after manual pollination (14 HAP) of emasculated flowers. Growth was visualized by aniline blue staining and length of pollen tubes and pistils were measured using ImageJ. *Wild-type* (n=17 pistils); *hda19-3* (n = 17 pistils); *hda19-3 STMRNAi* (pool of lines 1 and 10) (n=18 pistils). Letters over box-plots indicate statistical differences as determined by a one-way ANOVA test (p<0.05) and a Bonferroni *post hoc* test for multiple comparisons.

Ovule number and density increased in the *hda19-3 STMRNAi* plants according to the level of *STM* downregulation (Figure 5B), suggesting that *STM* expression level in the gynoecium was related to proper ovule number determination. Lines 4 and 9 showed the mildest reduction of *STM* levels (ca. 75% of *hda19-3* levels) (Figure 5B), and in accordance did not show significant changes in ovule number. Line 5 was the only exception, as it showed a reduction in *STM* levels similar to line 1 (ca. 35% reduction) (Figure 5A), but while Line 1 showed an increase in ovule number, Line 5 did not (Figure 5B). As expected, placenta length remained unchanged in all inspected lines (Figure 5C). Therefore, the increase in ovule number represents an increase in ovule density (Figure 5B-C).

Regarding the TT, we selected lines 1 and 10 for the inspection of TT phenotypes as they were comparable in terms of *STM* expression levels and ovule density. Although the septum of Line 1 and 10 was still partially unfused (Supplemental Figure 5A-C), the ridges were contacting more closely (Supplemental Figure 5B, C) and the subepidermal cells exhibited a *wild-type* like phenotype (Supplemental Figure 5A, C). In accordance, pollen tube growth-rate improved in the transgenic lines respect to *hda19-3* (Figure 5D, Supplemental Figure 5D-F). We hand-pollinated *wild-type*, *hda19-3* and *hda19-3 STMRNAi* pistils (lines 1 and 10) with *wild-type* pollen and 14 hours after pollination (14 HAP) we stained them with aniline blue (Figure 5D, Supplemental Figure 5D-F). We determined in a preliminary experiment that in *wild-type* pistils pollen tubes reach the bottom of the pistil in 16 hours. Therefore, at 14 HAP we would be able to precisely quantify the differences between each genotype. At 14 HAP, pollen tubes grew through 70% of *wild-type* pistils and 60% of *hda19-3 STMRNAi* pistils, a significant increase in respect to *hda19-3*, where pollen tubes barely grew 40% of the length of the pistil (Figure 5D, Supplemental Figure 5D-F). Overall, even if the anatomical changes in the TT are mild, they were sufficient to improve pollen tube growth in respect to *hda19-3* mutant.

In conclusion, our results strongly suggest that *STM* ectopic expression (Figure 4A-H) is mostly responsible for the ReT phenotypes present in *hda19*-3 (Figure 1) as reducing *STM* expression in *hda19-3* to *wild-type* levels leads to a recovery of ovule number and density (Figure 5B, C) and of TT function (Figure 5D, Supplemental Figure 5D-F).

### *stk* shows defects in the reproductive tract and increased histone acetylation at *STM* locus

HDA19 does not bind DNA directly, therefore, it has to be recruited to the DNA by transcription factors and other cofactors to regulate its targets. In the IM, HDA19 interacts with ETT, ARF4 and FIL (Chung et al., 2019b) to downregulate *STM*. However, in the CMM, the action of HDA19 is probably mediated by a different complex as at stages 8-9, ETT, ARF4 and FIL are mainly expressed in the lateral domains of the gynoecium (Sessions et al., 1997; Pekker et al., 2005; González-Reig et al., 2012).

Among the transcriptional factors expressed contextually to *HDA19* in the CMM, we highlighted STK as a possible candidate to partner with HDA19 to regulate *STM* expression. STK acts in redundancy with SHP1/2 to determine the ovule identity. Beyond this, STK is also required for septum and TT development (Colombo et al., 2010; Herrera-Ubaldo, et al., 2019; DiMarzo, et al., 2020), and displays repressor activity associated with histone deacetylation of its targets (Mizzotti et al 2014). To verify the role of STK, and potentially of SHP1/2, in the regulation of the expression of *STM* in the CMM, we analysed *STM* expression between *wild-type* and *stk shp1 shp2* mutant pistils by ISH (Figure 6A-H). *STM* resulted ectopically and overexpressed in *stk shp1 shp2* (Figure 6E-H), similarly to what we previously observed in *hda19-3* (Figure 4A-H). This observation suggests that STK and/or SHP1/2 could be involved in *STM* regulation during ReT development. Subsequentially, we measured ovule number and placenta length. We observed that *stk* mutant showed lower ovule number (Figure 6I) without changes in placenta length (Figure 6J), similarly to *hda19-3* (Fig. 1). Nevertheless, no statistically significative additive effects were noted in the *stk shp1 shp2* triple mutant (Fig. 6I, J), suggesting that SHP1 and SHP2 are not likely be involved in ovule number determination.

**Figure 6.**
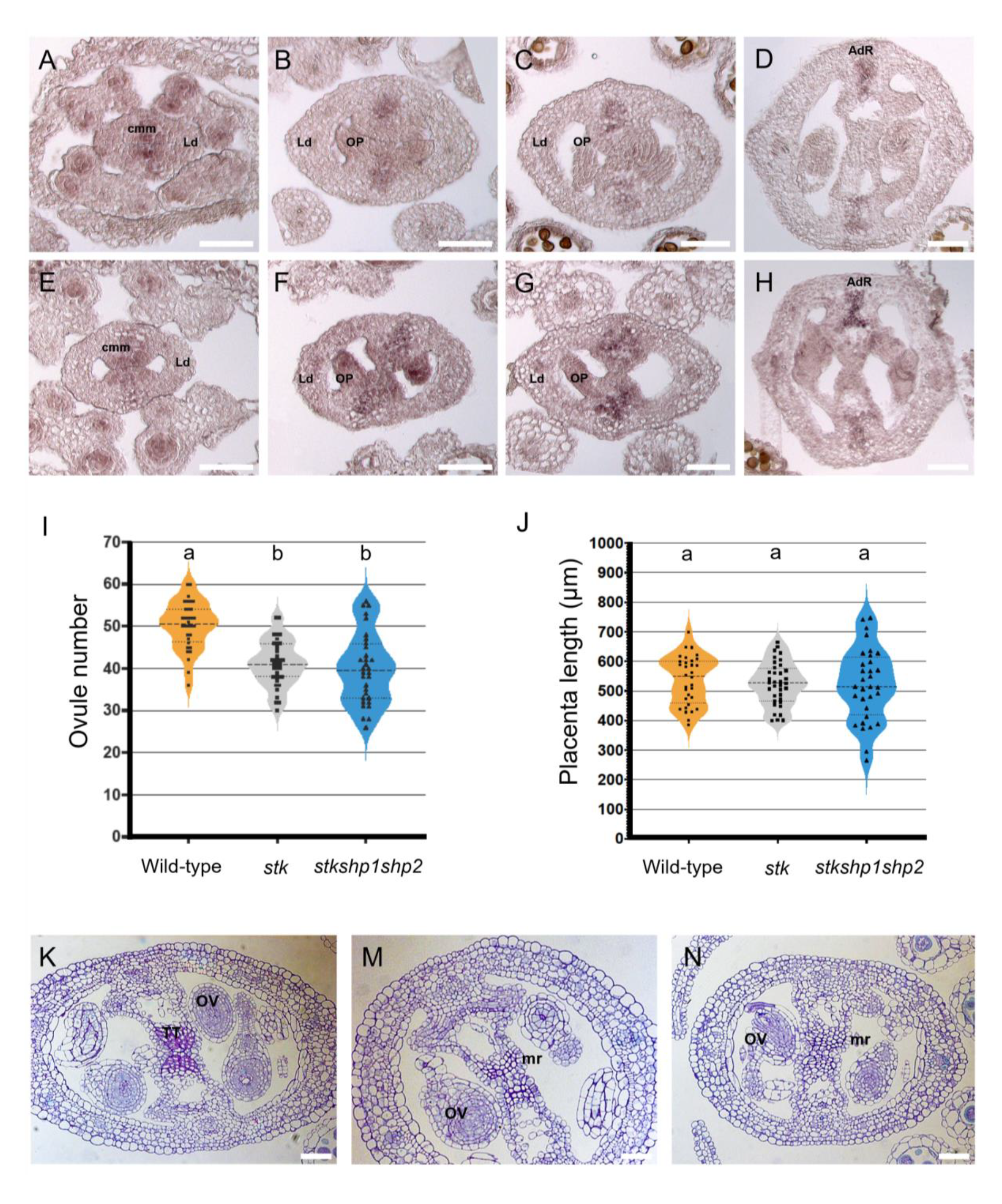
STK regulates *STM* expression in the CMM. **A-H)** Detection of *STM* mRNA in *wild-type* (**A-D**) and *stk shp1 shp2* mutant (**E-H**) by *in situ* hybridization. *STM* is over and ectopically expressed in *stk shp1 shp2* triple mutant at stages 7-10 of pistil development (**F-G**) in respect to the *wild-type* situation (**B-C**). Abbreviations: CMM = carpel margin meristem; LD= lateral domain; OP = ovule primordia; AdR = adaxial replum. Scale bar = 50 µm. **I-J)** Ovule number (**I**) and placenta length (**J**) in *wild-type, stk* and *stk shp1 shp2* mutants. Letters over violin-plots indicate statistical differences as determined by a one-way ANOVA test (p<0.05) and a Bonferroni *post-hoc* test. **K-N)** Transversal sections of stage 10-11 pistils of *wild-type* (**K**), *hda19-3* (**M**) and *stk* (**N**). *hda19-3* and *stk* show defective subepidermal cells in the TT. Abbreviations: TT = transmitting tract; OV = ovule; mr = medial ridge. Scale bar = 50 µm.

Finally, to corroborate our hypothesis that STK and HDA19 collaborate to downregulated *STM* to drive ReT differentiation, we evaluated the TT morphology in *stk* pistils. As expected, we found that subepidermal cells of *stk* ‘s medial ridges appear like those of *hda19-3* (Figure 6K-N).

### STK binds directly *STM* locus and is involved in its deacetylation

To explore whether STK activity was required to regulate *STM* acetylation state, we performed ChIP-qPCR using an anti AcH3K9/14 antibody on flowers from *wild-type*, *hda19-3, stk* and *stk shp1 shp2* (Figure 7A, B), testing the acetylation pattern of the regions described previously (Figure 4I). Similar to *hda19-3, stk* and *stk shp1 shp2*, showed an increase in H3K9/14 acetylation at the level of the TSS, and a moderate increase at the promoter (Figure 7A, B), suggesting a similar function of HDA19 and STK in regulating the acetylation state of the regions tested.

**Figure 7.**
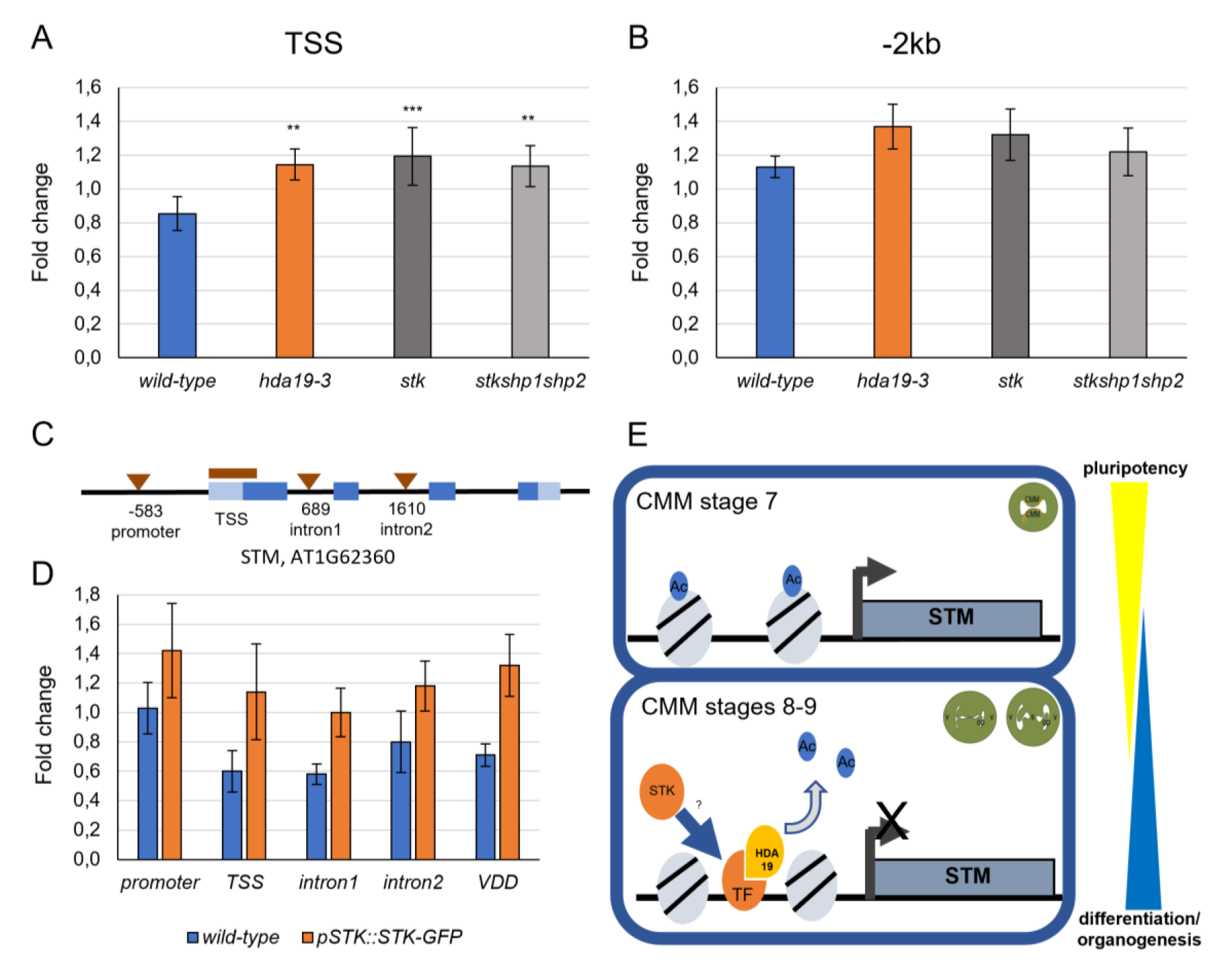
STK binds to *STM* locus. **A-B)** AcH3K9 ChIP-qPCR of *STM* TSS (**A**) and of a distal promoter region (−2kb) (**B**) in *wild-type*, *hda19-3*, *stk* and *stk shp1 shp2* mutants. The pattern of acetylation in the *STM* locus is similar among the different mutant genotypes. Graphs show the mean ± sd value of enrichment among three independent biological replicates of the AcH3K9 ChIP-qPCR experiment performed on *wild-type*, *hda19-3, stk and stkshp1sp2* flowers. Student’s T-test, ** = p-value < 0. 01, *** = p-value < 0. 001. **C)** Regions of *STM* locus tested by anti-GFP ChIP-qPCR on *wild-type* and *pSTK::STK-GFP* flowers. Triangles mark CaRG boxes detected by AthaMap (Steffens et al., 2004). The differentially acetylated region at the TSS of *STM* is marked with an orange line. **D)** ChIP-qPCR to test binding of STK (*pSTK::STK-GFP*) to the promoter, TSS, first intron and second intron of *STM*. *VDD* is a known direct target of STK, so it was used as positive control. Graph shows the mean ± sd enrichment value among four independent biological replicates of the anti-GFP ChIP-qPCR experiment. **E)** Model of the regulation of *STM* expression in the CMM involving HDA19 and STK. Until stage 7, the expression of *STM* in the CMM would be high thanks to the presence of acetylated histones in association with *STM* locus. At stages 8-9, when the CMM has to start producing the reproductive tract organs, HDA19 would deacetylate the histones associated with *STM* locus, to downregulate it and allow the formation of ovule primordia and TT. Direct binding of STK to *STM* locus would facilitate the action of HDA19, potentially by generating multimeric complexes involved in HDA19 binding and stabilization to *STM*.

Finally, we verified whether STK could directly bind to the *STM* genomic region by performing a ChIP-qPCR using an anti-GFP antibody on flowers of *wild-type* and *pSTK::STK-GFP* plants. MADS-box proteins bind DNA motifs called CarG boxes. We found, and tested by ChIP-qPCR, CArG boxes in the proximal promoter area, in the first intron and second intron of *STM* (Figure 7C). In addition, we also tested the region near the TSS of *STM* as it was the most enriched in the AcH3K9/14 ChIP-qPCR (Figure 7A). As a positive control we used *VERDANDI* (*VDD*), a direct target of STK (Matias-Hernandez et al., 2010) (Figure 7D). The first *STM* intron showed the strongest enrichment, followed by the TSS. The second intron and the promoter showed no enrichment (Figure 7D). This result indicates that STK can directly bind the *STM* locus, and that its action is required to regulate its expression level (Figure 6 A-H).

Based on our phenotypic analysis, expression studies and ChIP-qPCR experiments, we show that HDA19 and STK have a similar effect over *STM* in the regulation of its levels and domain of expression. Such activity is required to regulate the CMM functionality and allow ReT development and ovule number determination, both traits essential for plant fitness (Figure 7E).

## DISCUSSION

In this work, we observed that HDA19 is necessary for ovule initiation and TT differentiation (Figure 1, Supplemental Figure 1). To study the cause of these defects, we performed an RNA-seq on ReT cells sorted by FACS with the aid of the *pSTK::STK-GFP* line. The transcriptomic analysis has given an overview of gene expression in tissues expressing STK-GFP fusion protein that include the CMM, TT, placenta and late sporophytic ovule tissues. Comparing cells expressing STK-GFP in *wild-type* and *hda19-3* allowed us to focus on the role of HDA19 in these tissues. Consistent with *hda19-3* phenotype, several genes involved in meristem maintenance, including *STM,* were upregulated in *hda19-3* ReT. Instead, genes involved in organogenesis like *CUC1/2*, *ANT*, *PAN*, *PIN1,* or *HEC1* and *NTT* were downregulated in *hda19-3* sorted cells (Figure 3). Our results show that HDA19 is required to repress *STM* in the CMM to allow organ formation and tissue differentiation, and that STK is also involved in this process. In plants in which either *HDA19* or *STK* are mutated, the *STM* locus is more acetylated respect to the *wild-type* situation. This leads to *STM* upregulation and ectopic expression in the CMM leading to a reduced ovule number and defects in TT differentiation.

To assess whether *STM* de-regulation was the main responsible for the ReT phenotypes of *hda19-3*, we reduced *STM* levels in *hda19-3* plants. *stm* loss of function mutants lack a functional SAM and consequently fail to produce flowers. Therefore, we used a weak constitutive promoter, showing that the downregulation of *STM* in *hda19-3* improved ovule number and density and growth rate of pollen tubes in the TT (Figure 5, Supplemental Figure 5). None of the transgenic lines obtained rescue completely the *hda19-3* phenotype. This is not surprising as HDA19 regulates many processes, so it is unlikely that ectopic expression of *STM* is the only cause of all *hda19-3* defects. In addition, *KNAT6* and *BP*, which are partially redundant with STM in the SAM (Byrne et al., 2002) were also upregulated in *hda19-3*’s CMM, therefore, their downregulation might be necessary for a better complementation.

Previously, Scofield and collaborators observed that the overexpression of *STM* using an inducible overexpression system led to the transformation of ovules into carpelloid structures (Scofield et al., 2007). This aligns with our observations but represents a more extreme phenotype probably because the levels of *STM* expressed in the flower of the transgenic plants were much higher than those observed in *hda19-3.* If STM negatively regulates *STK, SHP1* and *SHP2*, its ectopic expression in the ovules could lead to a downregulation of STK and SHP1/2 strong enough to cause the transformation of the ovule integuments into carpelloid structures as described by Scofield et al., 2007. Our transcriptome analysis clearly showed the downregulation of genes necessary for organ primordia formation as well as ovule identity in *hda19-3* (Figure 3), which might be due to the action of STM. This is similar to what happens in other meristems like the SAM or the IM (Long et al., 2006; Chung et al., 2019) providing functional evidence to support the idea that the CMM functions in a similar way as its indeterminate counterparts. However, how its activity is terminated requires further research, as ISH showed that after stage 10, *STM* expression in *hda19-3* is similar than in *wild-type* (Figure 4A-I), suggesting that besides HDA19, other factors contribute to *STM* silencing in the CMM, since lack of HDA19 delays *STM* silencing but does not avoid it completely.

*STK* was first described as a member of the ovule identity complex redundant with *SHP1/2* (Pinyopich et al., 2003; Favaro et al., 2003). However, STK and SHP1/2 have also non-redundant functions along reproductive development (Liljegren et al., 2000; Pinyopich et al., 2003; Mizzotti et al., 2012, 2014, Ezquer et al., 2016; DiMarzo et al.,2020). Our results support that STK is the main player required for HDA19’s action over *STM* in the CMM as the pattern of histone acetylation of the *STM* locus is similar in both mutants (Figure 7A-B). However, while *stk* shows a reduction of ovule number and density similar to *hda19-3*, the phenotype regarding the TT in *stk* is milder than *hda19-3*’s. The role of STK in the development of the TT was already described (Marsch-Martínez et al., 2014; Herrera-Ubaldo et al., 2019, Di Marzo et al., 2020), but single *stk* mutants showed very mild defects that became more severe when *stk* was combined with *ntt* and *ces* mutants. Overall, this suggests that in the TT, HDA19 might be partnering with other proteins besides STK to regulate *STM*.

Finally, we found that STK binds *STM* locus at the level of the TSS and the first intron (Figure 7C-D). MADS-box proteins form tetramers able to bind simultaneously to two DNA binding sites, creating DNA loops (Smaczniak et al., 2012). As the TSS is the region that is over-acetylated in *hda19-3* and *stk* mutants, and STK binds at the level of the TSS and the first intron, a plausible model could be that a MADS-box complex containing STK and HDA19, binds to *STM* locus forming a loop between the TSS and the first intron. Then, HDA19 would deacetylate the TSS region of *STM* reducing its expression, allowing organ initiation from the CMM (Figure 7E). It has been previously shown that STK displays both activator and repressor properties (Mizzotti et al., 2014), suggesting that only a fraction of its direct targets are regulated through histone de-acetylation (Mizzotti et al., 2014). This supports STK ability to work in different multimeric complexes involving different partners. By Yeast Two Hybrid assay we have not been able to highlight a direct interaction between HDA19 and STK. For this reason, it is likely the STK and HDA19 interaction is indirect and/or regulated by post-translational modifications. Exploring this is outside the scope of the present work, but it will be extremely interesting to investigate the mechanism of their interaction in the future.

Another interesting observation is that *STK* expression starts at stage 8, in the flat placenta, well before OP start to form. In light of our results, the onset of *STK* expression at stage 8, would lead to the reduction of the levels of *STM*, allowing ovule initiation and septum fusion (Pinyopich et al., 2003). This idea is supported by the observation that the levels of *STM* are normal in *hda19-3* before stage 8 (Figure 4 A-H) suggesting that HDA19 exerts its action over *STM* during stages 8-10, when STK starts to accumulate.

To our knowledge, the only work on transcriptomics of sorted gynoecium cells was done with the aid of a dexamethasone-inducible *pAP1::AP1-GR ap1cal* system (Villarino et al., 2016). This system allows to obtain many synchronized flowers, but dexamethasone can introduce variability and/or artifacts. Indeed, the authors observed occasional ectopic expression of SHP2-YFP outside the normal expression domain of *SHP2* (Villarino et al., 2016). Our work demonstrates that it is possible to perform consistent FACS-coupled transcriptomics of gynoecium cells using markers of moderate expression without the aid of inducible systems. Moreover, as our dataset contains TT cells and placenta and sporophytic ovule cells, it will be highly useful for future studies of CMM and ovule development. Indeed, we already found hints of the possible role of several genes (Wynn et al., 2011; Villarino et al., 2016). For example, *YAB3* was proposed to have a non-cell autonomous role in CMM development (Wynn et al., 2011), but our data indicates otherwise (Figure 3). *YAB3* represses medial factors (González-Reig et al., 2012) and its mutation affects CMM development (Nole-Wilson, 2006; Wynn et al., 2011b). Wynn and collaborators did not detect *YAB3* in the medial domain by ISH, so its effect was proposed to be non-cell autonomous (Wynn et al., 2011). As *pSTK::STK-GFP* is not expressed in the lateral domain, our data indicates that *YAB3* is indeed expressed in the middle domain at low levels in the wild-type that might be undetectable by ISH.

Our results show that the tight control of *STM* levels achieved through the action of HDA19 and STK are essential to maximize reproductive fitness of the plant, as even moderate changes in its expression lead to defects in the ReT that reduce seed set.

## EXPERIMENTAL PROCEDURS

### Plant materials and growth conditions

Plants for phenotypical analysis were grown in a greenhouse under long day conditions (16 hours light, 8 hours dark) at 22-24 °C.

For sorting experiments, plants were grown in a walk-in growth chamber under continuous light at 22 °C.

Wild-type Col-0 seeds were already available in the laboratory. *hda19-3* (SALK_139445) was provided by the Nottingham Arabidopsis Stock Center (NASC).

*pSTK::STK-GFP* line was available in the laboratory and first described in Mizzotti *et al*., 2014.

*Stk* and *STK +/- shp1 shp2* seeds (used to obtain the *stk shp1 shp2* triple mutant from segregation) were already available in the laboratory and previously described by (Pinyopich et al., 2003; Favaro et al., 2003).

### Ovule number and placenta length measurement

Inflorescences were fixed overnight in ethanol:acetic acid 9:1 and rehydrated in an ethanol series. Pistils (stages 6-10) were dissected and mounted on chloral hydrate:glycerol:water (8:1:3, w/v/v) and immediately observed under a Zeiss Axiophot D1 microscope (Carl Zeiss MicroImaging) equipped with DIC optics and an Axiocam MRc5 camera (Zeiss) with Axiovision software (version 4.1). Ovules were counted manually. Placenta length was measured using Axiovision software.

### Pistil histology and imaging

Individual flowers from *wild-type*, *hda19-3, 35sRNAi:STM* and *stk* were fixed in 2% (w/v) paraformaldehyde and 2,5% (w/v) glutaraldehyde in PIPES buffer [0.025 M, pH 7, 0,001 % (v/v) Tween-80], placed under vacuum for 1h and then at 4°C overnight. Material was dehydrated in an ethanol series and embedded in LR White resin. Thick sections (500 µm) were obtained with a Leica EM UC7 Ultramicrotome stained with 1 % (w/v) toluidine blue (Sigma-Aldrich, St Louis, MO, US) and mounted with DPX (Sigma). Slides were observed under a Zeiss AxioImager AZ microscope equipped with a Zeiss Axiocam MRc3 camera using Zen Imaging software (Zen 2011 SP1).

### Aniline blue staining for pollen tube growth analysis

Flowers were emasculated and pollinated with *wild-type* pollen. Pollen tubes were allowed to grow for different times depending on the experiment (12, 14 or 24h). Then, pistils were fixed overnight in 9:1 ethanol:acetic acid, washed three times with water, incubated in 1M NaOH overnight, washed three times in water and incubated in 0,1% aniline blue (w/v). Pistils were imaged using a Zeiss AxioPhot D1 microscope equipped with fluorescence filters. Images were recorded with an Axiocam MRc5 camera (Zeiss) with Axiovision software (version 4.1). Pollen tubes length was measured using ImageJ (Schneider et al., 2012).

### Protoplast recovery and Fluorescence Activated Cell Sorting (FACS)

*pSTK::STK-GFP* construct was introgressed into *hda19-3* background and then used for cell sorting, along with the parental *pSTK::STK-GFP*. Starting material consisted in flowers from stages 1-12. Protoplasting and sorting was performed as described by Villarino et al., 2016, with some modifications. Cells were sorted into RLT buffer from the RNAeasy Microkit (Qiagen) supplemented with 1% β- mercaptoethanol (v/v), keeping a ratio of 3 volumes of RLT buffer per 1 volume of sorted cells (in our sorting conditions 50,000 cells occupied 200 µL). Cell sorting was performed using a Moflo XDP (Beckman Coulter Inc.). Each experiment yielded between 80,000 and 100,000 GFP-positive cells.

### RNA extraction and expression analysis

RNA was extracted immediately after sorting using an RNAeasy Microkit (Qiagen) with a few modifications. We used one column for every 25,000 cells. Columns were eluted twice, using the first eluate for the second elution. Finally, all eluates from the same sample were pooled, yielding 70-100 ng of total RNA. For qRT-PCR validation, the columns were eluted a third time, using 14 µL of nuclease-free water that was sequentially used to elute all the columns corresponding to the same sample. This yielded around 5 ng of total RNA that was retro-transcribed using SuperScript III First-Strand Synthesis System (Invitrogen/Life Technologies). Then, expression of *STK-GFP* was measured by qRT-PCR using a Thermal Cycler from Applied Biosystems and a QuantiTect SYBR Green PCR Kit (Qiagen). *MON1* (*AT2G28390*) was used as a reference gene for normalization based on Czechowski et al., 2005.

For evaluation of *STM* levels in RNAi lines, RNA was extracted using Trizol (Invitrogen) and 500 ng of RNA were retrotranscribed using iScript cDNA synthesis kit (BioRad). qRT-PCRs were performed using iTaq Universal SYBR Green Supermix (BioRad) in a Bio-Rad iCycler iQ thermal cycler. *ACT8* gene (*AT1G49240*) was used as reference gene for normalization because, according to our RNA-seq data, its RNA levels do not change in *hda19-3* plants.

All primers used in qRT-PCR experiments showed an amplification efficiency close to 100%, so the 2^−ΔΔCT^ method was used for calculation of expression changes.

All primers used are listed in Supplemental Table 1.

### RNA sequencing and analysis

Three validated samples of RNA from *wild-type* GFP-positive cells and 4 samples from *hda19-3* GFP-positive cells were used for library preparation. Libraries were sequenced in one lane of the HiSeq2500 Illumina platform with a yield of 250 million reads and an average of 35 million reads per sample. Around 10 million reads were filtered out and of the 240 million reads remaining, 230 million mapped uniquely (96% of the reads after filtering) and were used for subsequent analysis.

All steps of library preparation, RNA sequencing and bioinformatics analysis were essentially performed as in Villarino et al., 2016. Briefly, we used Cufflinks (Trapnell et al., 2012), DeSeq2 (Love et al., 2014) and EdgeR (Robinson et al., 2009) to analyze the RNA-seq data (Supplemental File 1-3). Then, we crossed the data obtained by the 3 software and selected the DEGs detected by the three programs, with an FDR < 0.001 and with a logFC > |1.2| (Supplemental File 4). Genes whose expression value was zero in either *wild-type* or *hda19-3* were filtered out.

### Chromatin immunoprecipitation and qPCR analysis

Chromatin immunoprecipitation was performed as described in Mizzotti et al., 2014, using a rabbit anti-H3 acetyl K9 (Upstate 07-352, Sigma-Aldrich) and a mouse monoclonal anti-GFP antibody from Roche (11814460001).

qPCR was performed using iTaq Universal SYBR Green Supermix (BioRad) in a Bio-Rad iCycler iQ thermal cycler.

Enrichment of tested areas was calculated as fold change against a control non-enriched region (*GAPDH* locus) as previously described by Matias-Hernandez et al., 2010. Primers used for ChIP-qPCR are listed in Supplemental Table 1.

### *In-situ* hybridization

Sectioning and *in situ* hybridization were performed as described by (Dreni et al., 2007). *STM* probe was obtained as described in Simonini & Kater, 2014.

### Generation of RNAi lines

For *35s::STM-RNAi* construct, a fragment of 352 bp of *STM* coding sequence was amplified (see primer list in Supplemental Table 3), cloned into pDONR207 and then subcloned into pFRH (derived from pFGC5941; NCBI accession number AY310901) using Gateway technology (Invitrogen).

The construct was transformed into *Agrobacterium tumefaciens* strain EHA105 and *hda19-3* plants were transformed by floral dip (Clough and Bent, 1998). Transformants were selected in MS media (Murashige and Skoog, 1962) supplemented with 20 mg/L of hygromycin. Presence of the construct was confirmed by genotyping (see primers in Supplemental Table 1).

### Clustering analysis

Heatmaps were obtained using Heatmapper (Babicki et al., 2016). Genes were hierarchically clustered by Euclidean distances using average linkage method.

### Data availability

RNA-seq data are available at the accession number (XXXXX)

### Funding

Manrique S, Cavalleri A, Colombo L, were founded by MIUR PRIN2017. Guazzotti A, Colombo L were founded by H2020-EU.1.3. (SEXSEED ID: 690946). Manrique S was founded by EMBO SHORT TERM (2015). Manrique S, Masiero S, Mizzotti C, were founded by MC-IRSES - International research staff exchange scheme (IRSES) (FRUITLOOK ID: 612640). Coimbra S was founded by SeedWheels FCT Project - POCI-01-0145-FEDER-027839.

### Author contribution

Conceived and design the research: MS, CL, CA, GA, FR. Performed the experiments: MS, CA, GA, VG, OE, PAM, CS, SS. Contributed reagents/materials/analysis tools: CL, FR, HT, GU. Analysed the data: MS, AC, VG, BA. Wrote the manuscript: MS, CA, CL, GA

## Abbreviations

CMM: Carpel margin meristem
TT: transmitting tract
OP: ovule primordia
ReT: reproductive tract
ISH: *in situ* hybridization
ChIP: chromatin immunoprecipitation

## Acknowledgments

Thanks to Sarah Schuett (NCSU) and Claudia Bazzini (UniMi) for technical assistance with FACS and the Genomic Sciences Laboratory Research Facility (NCSU) for library preparation and Illumina sequencing.

