## Supplemental Figure and Tables for "HISTONE DEACETYLASE 19 REGULATES *SHOOT MERISTEMLESS* EXPRESSION IN THE CARPEL MARGIN MERISTEM CONTRIBUTING TO OVULE NUMBER DETERMINATION AND TRANSMITTING TRACT DIFFERENTIATION"

### Supplemental Figures

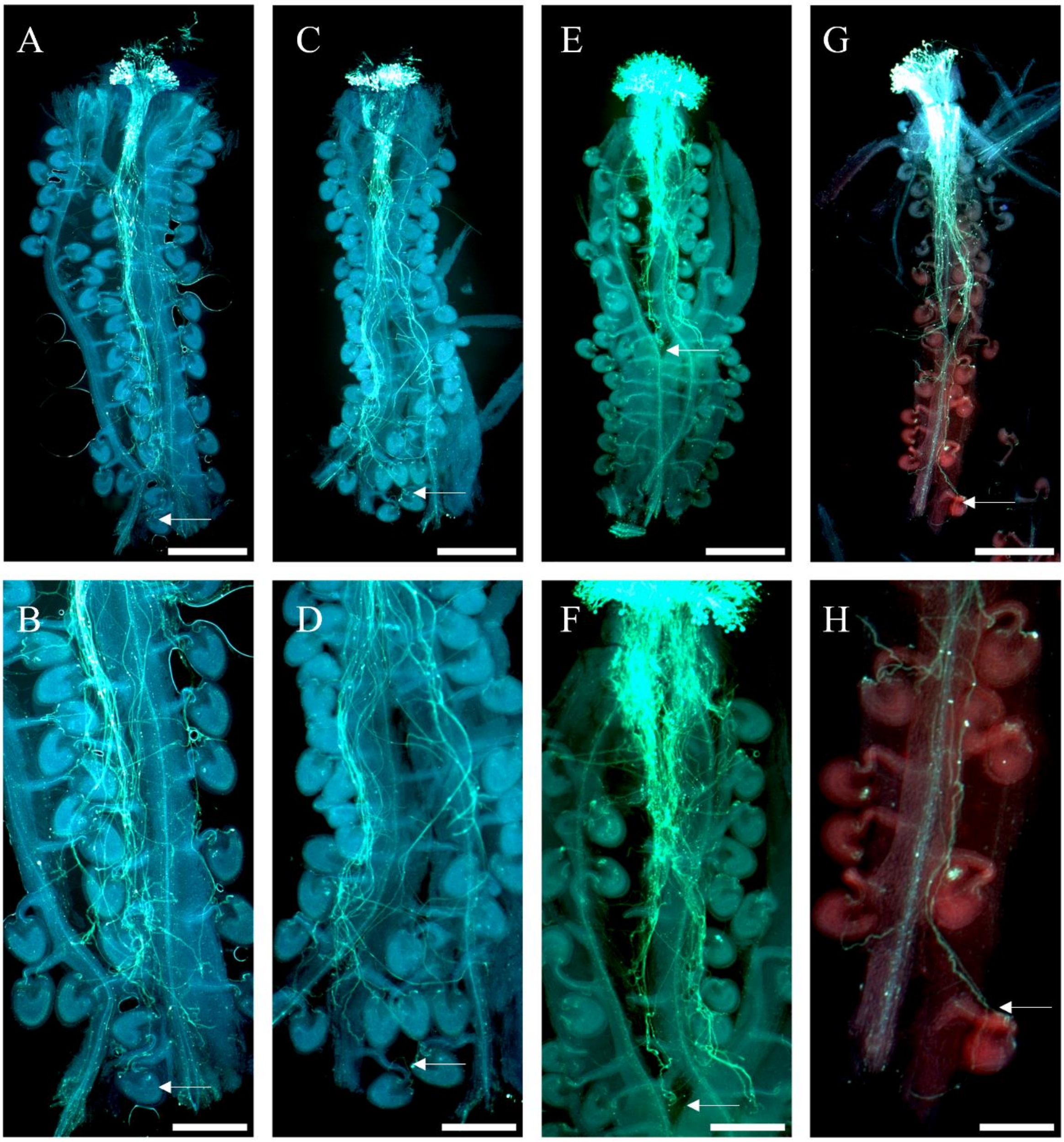


**Supplemental Figure 1. Growth of pollen tubes in pistils of *wild-type*, *hda19-3* +/- and *hda19-3* -/- plants. A-B)** pollen tubes in a *wild-type* pistil at 12 HAP. **C-D)** pollen tubes in an *hda19-3* heterozygous pistil at 12 HAP. **E-F)** Pollen tubes in an *hda19-3* homozygous pistil at 12 HAP. **G-H)** Pollen tubes in an *hda19-3* homozygous pistil at 24 HAP. Arrowheads mark the tip of pollen tubes. Images **B**, **D**, **F** and **H** (Scale bars = 50 µm) are enlargements of the portion of the pistil containing the longest pollen tubes of images **A**, **C**, **E** and **G** respectively (Scale bars = 100 µm).

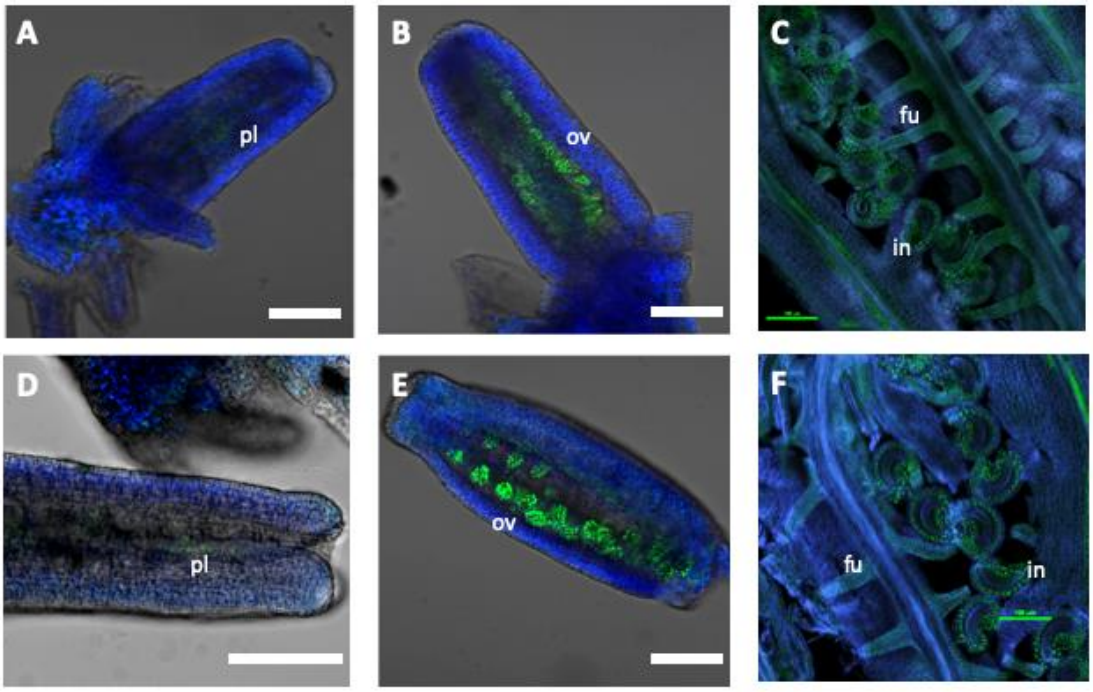


**Supplemental Figure 2. Expression of *pSTK::STK-GFP* marker line. A-C)** *wild-type* pistils at stages 8 (**A**), 9 (**B**) and 11 (**C**). **D-F)** *hda19-3* pistils at stages 8 (**D**), 9 (**E**) and 11 (**F**). The spatial expression pattern of the *pSTK::STK-GFP* marker line is maintained in both genotypes. First hints of expression are observed at stage 8 in the flat placenta (**A**, **D**). At stage 9, a strong expression in the ovule primordia and in the placenta can be observed, with weaker expression in the transmitting tract (**B**,**E**). At stage 11, fluorescence appears in the ovule integuments, funiculus and placenta (**C**, **F**). Images were obtained with an A1 Nikon confocal microscope. Abbreviations: pl=placenta; OV= ovule; fu= funiculus; in= integuments. Scale bar = 20 nm.

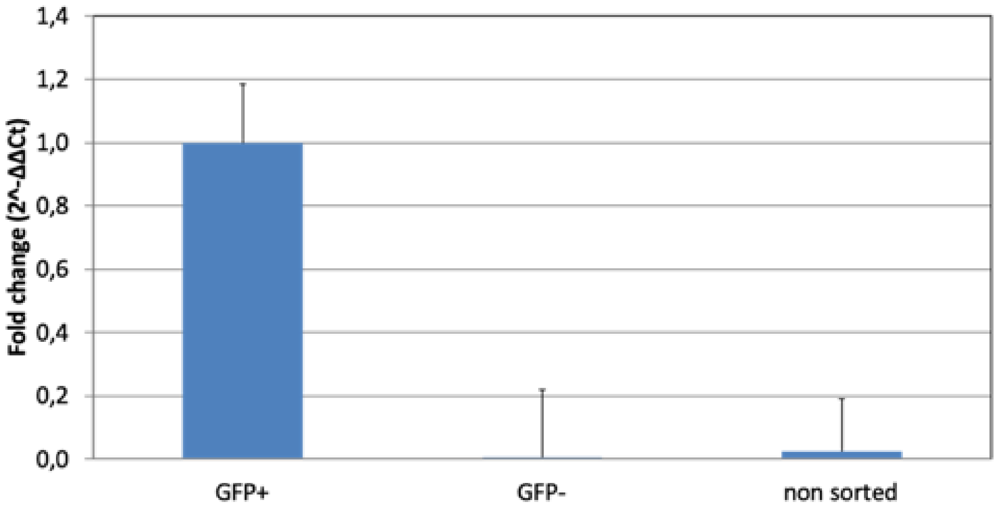


**Supplemental Figure 3.** Enrichment of *STK-GFP* transcript in cells sorted as GFP+, GFP- and non-sorted. *MON1* (*AT2G28390*) was used as reference gene for normalization.

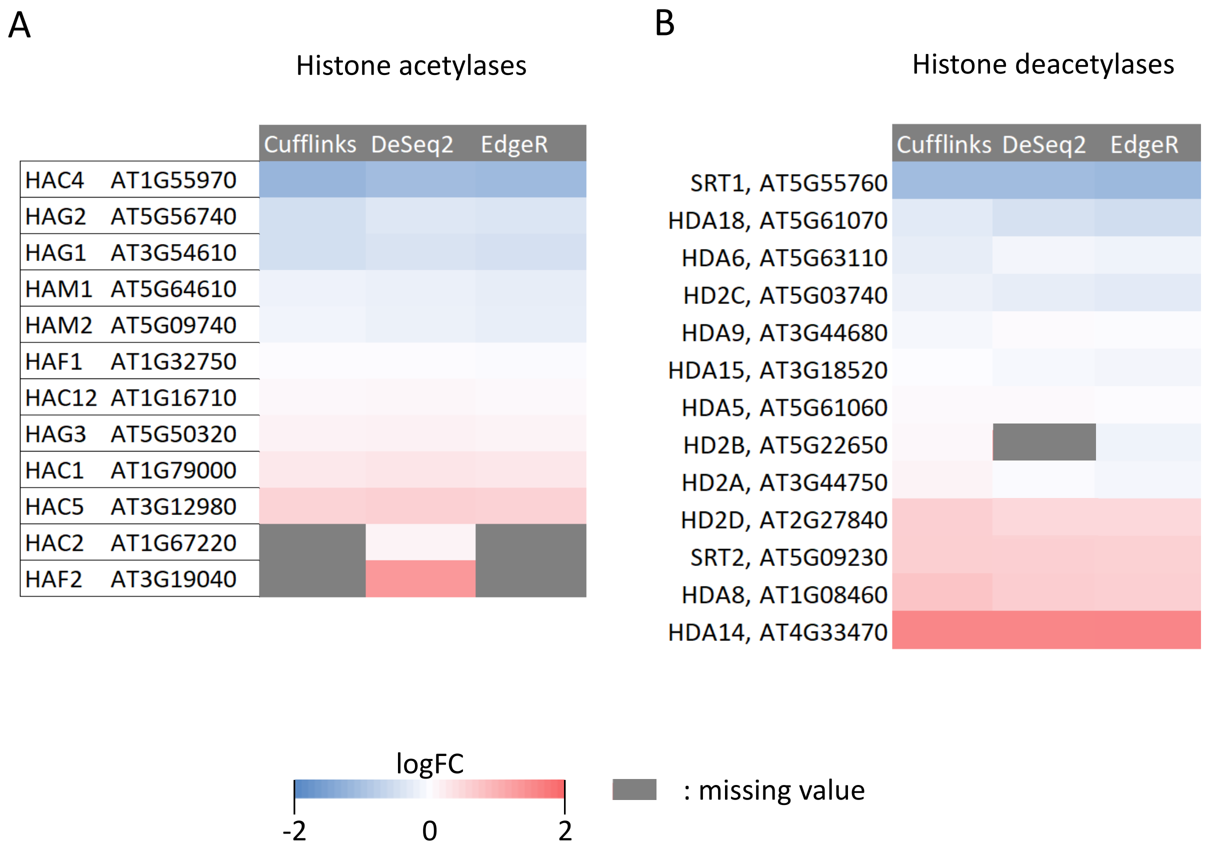


**Supplemental Figure 4. Expression levels of Arabidopsis histone acetylases and deacetylases in *hda19-3 pSTK::STK-GFP* expressing cells compared with the wild-type. A)** Expression of histone acetylases in *hda19-3 pSTK::STK-GFP* expressing cells according to Cufflinks, DeSeq2 and EdgeR. HAF2 is somewhat upregulated according to DeSeq2, but filtered out by Cufflinks and EdgeR, suggesting its expression pattern was inconsistent in our samples **B)** Expression of histone deacetylases in *hda19-3 pSTK::STK-GFP* expressing cells according to Cufflinks, DeSeq2 and EdgeR.

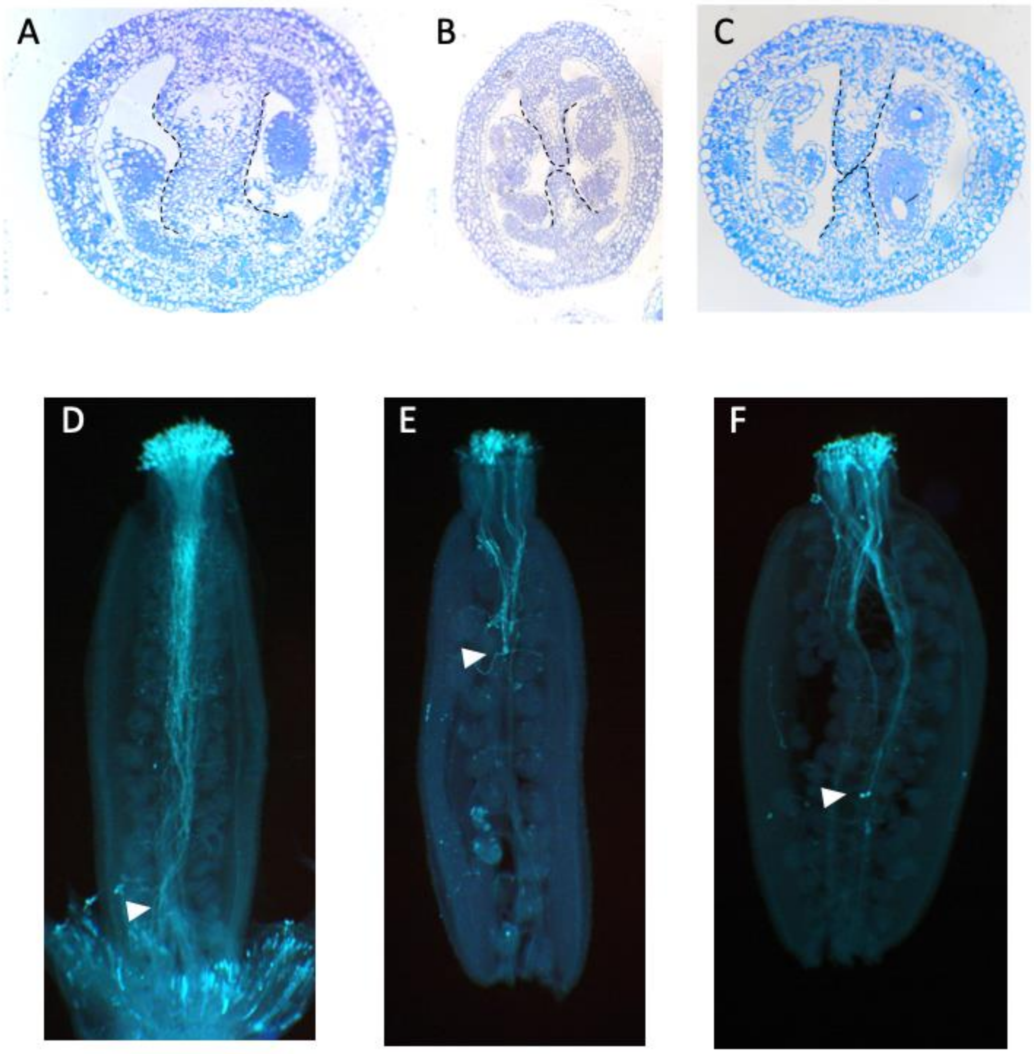


**Supplemental Figure 5. Transmitting tract phenotypes of *wild-type*, *hda19-3* and *hda19-3 STMRNAi* plants**. **A-C)** Transversal sections of *wild-type* (**A**), *hda19-3* (**B**) and an example of an *hda19-3 STMRNAi* line (**C**). **D-F)** Aniline blue staining of pollen tubes at 14 HAP in *wild-type* (**D**), *hda19-3* (**E**) and one example of *hda19-3 STMRNAi* (**F**). White arrowheads indicate the position of pollen tubes. Scale bar = 20 µm.

### Supplemental Tables

#### Supplemental Table 1: Primers used in this study

| Primer | Sequence | Usage |
|---|---|---|
| STK-GFP FWD | TTGAGCTTGACAATGAGAACATC | Expression analysis of *STK* gene (validation of sorted RNA samples) |
| STK-GFP REV | GGTTTTTCGCAGACGAGATT | |
| MON1 FWD | CAGACAAGGCGATGGCGATA | Expression analysis of *MON1* gene |
| MON1 REV | GCTTTCTCTCAAGGGTTTCTGGGT | |
| ACT 8 FWD | CTCAGGTATTGCAGACCGTATGAG | Expression analysis of *ACT8* gene |
| ACT8 REV | CTGGACCTGCTTCATCATACTCTG | |
| STM ISH FWD | GTTGCTTCTTCTTCTTCTCC | Synthesis of STM probe for *in situ* hybridisation |
| STM ISH REV + T7 | TAATACGACTCACTATAGGGACGAGCATTTCACAGTAAGC | |
| STM_FWD | CCTTCAACGTGTCGAGTGTC | Expression analysis of *STM* gene |
| STM_REV | ACTTCTTCCTCGGATGACCC | |
| STM RNAI FWD | GGGGACAAGTTTGTACAAAAAAGCAGGCTAACCCTTGCTCCTCTTCCTC | Gateway cloning of a fragment of *STM* CDS for RNAi construct |
| STM RNAI REV | GGGGACCACTTTGTACAAGAAAGCTGGGTACCGGAGAAAGAGGAAGGTG | |
| STM_RNAI_GTPYING FWD | CCTCTGTCAAGGCCAAGATC | Genotyping of STM-RNAi plants (construct is distinguished from endogenous gene because primers span two exons) |
| STM_RNAI_GTPYING REV | GACACTCGACACGTTGAAGG | |
| STM_H3K9_CHIP_FWD | ACTTTGTTGGTGGTGTGACTG | *STM* TSS region |
| STM_H3K9_CHIP_REV | ATGATGATGATGATGCCGCC | |
| STM_H3K9_CHIP_FWD | TCTATGAGCGTAGGAGAC | *STM* -2Kb region |
| STM_H3K9_CHIP_REV | CCAAAATATGTTGGATCTGGAC | |
| STM_CARG_I1_FWD | TGTTTACTAGTTACTTAACCCAGCT | *STM* first intron CARG-BOX |

| | | |
|---|---|---|
| STM_CARG_I1_REV | TGGATAATCTCTTGCAAGTAGGGT | |
| STM_CARG_I2_FWD | TGCTCGTCCTTAGATCTATTGCT | *STM* second intron CARG-BOX |
| STM_CARG_I2_REV | AGTAGATGTGAGTTTGTGTGTCT | |
| STM_CARG_PROM_FWD | TTCACTGGACTTTCCGAGGC | *STM* promoter CARG-BOX |
| STM_CARG_PROM_REV | TGATTTCTCATTTATGCCTTTTCGGA | |
| VDD FWD | GGAAATATGACGCTTGTCTTTTTAG | Positive control STK-GFP CHIP |
| VDD REV | CAGAAACAGCAATATGCTCGTG | |
| GAPDH FWD | CTCGTTGTGCAGGTCTCAAA | Normalizer for CHIP experiments |
| GAPDH REV | CTAGTGGCTCATCGCAGA | |

**Supplemental Files**

**Supplemental File 1: Results of RNA-seq analysis with Cufflinks**

**Supplemental File 2: Results of RNA-seq analysis with DeSeq2**

**Supplemental File 3: Results of RNA-seq analysis with EdgeR**

**Supplemental File 4: Common DEGs among Cufflinks, DeSeq2 and EdgeR**

**Supplemental File 5: enrichment analysis of ‘Biological Process’ GO terms associated with the upregulated genes**

**Supplemental File 6: enrichment analysis of ‘Biological Process’ GO terms associated with downregulated genes**
